# The branched receptor binding complex of *Ackermannviridae* phages promotes adaptative host recognition

**DOI:** 10.1101/2024.03.21.586117

**Authors:** Anders Nørgaard Sørensen, Cedric Woudstra, Dorottya Kalmar, Jorien Poppeliers, Rob Lavigne, Martine Camilla Holst Sørensen, Lone Brøndsted

## Abstract

Bacteriophages may express multiple receptor binding proteins, enabling the recognition of distinct and diverse bacterial receptors for infection of a broad range of strains. *Ackermannviridae* phages recognize diverse O-antigens or K-antigens as receptors by expressing multiple tail spike proteins (TSPs). These TSPs interact and form a branched protein complex protruding from the baseplate attached to the distal tail. Here, we aimed to mimic the evolution of the TSP complex by studying the acquisition of new TSPs without disrupting the functionality of the complex. Using kuttervirus phage S117 as a backbone, we demonstrated the acquisition of entire *tsp* genes from *Kuttervirus* and *Agtrevirus* phages within the *Ackermannviridae* family. A fifth TSP was designed to interact with the complex and provide new host recognition to expand the branched TSP complex. Interestingly, the acquisition of *tsp5* resulted in new variants of the branched TSP complex due to the exchange or deletion of *tsp* genes. Overall, our study provides novel insight into the development of the branched TSP complex, enabling *Ackermannviridae* phages to adapt to new hosts.

## Introduction

Bacteriophages (phages) express receptor binding proteins that form long or short tail fibers (TF) or tail spike proteins (TSPs) attached as distal structures to the phage tail. When infecting a bacterial host, the phage receptor binding proteins recognize the surface receptor of the bacterial host. The specificity of this initial phage-host interaction ensures the infection of proper hosts, and receptors have been suggested as the major determinant of the host range of phages ^1^. On the phage side, most characterized phages typically use a single receptor binding protein to recognize their bacterial host. Yet, the rise of whole genome sequencing has identified an increasing number of phages that encode multiple receptor binding proteins. For example, gamaleyavirus phage G7C expresses two tail spike proteins (TSPs) interacting in a complex where one of the TSPs deacetylates the O-antigen on *E. coli* ^2^. Even more complex is the network of receptor binding proteins of recently characterized phages; for instance, phage LKp24 encodes 14 TSPs, each proposed to recognize a distinct K-antigen of *Klebsiella*, including K2, K1, K25, K35, and KN4 ^3^.

The multiple receptor binding proteins of these phages often form a branched complex by interacting through defined structural domains or entire proteins conserved even across phage families ^4,5^. For example, TSPs of *Klebsiella* phages carrying a branched TSP complex share similarities to domains 2 and 3 found in Gp10 of the T4 phage ^3,4^. In phage T4, Gp10 is part of the peripheral baseplate and consists of four domains that form an X shape and are crucial for assembling the tail fiber complex. Domains 2 and 3 of Gp10 adopt a conserved ß-jellyroll fold interacting with the short tail fiber and the baseplate protein Gp9, which further interacts with the long tail fiber ^6,7^. Therefore, the ability to mediate protein-protein interactions may explain why Gp10-like domains are widespread in phages expressing multiple receptor binding proteins arranged in a branched complex.

Phages of the *Ackermannviridae* family are another example of multiple TSPs linked together in a branched complex using Gp10-like domains. In these phages, Gp10-like domains are found as N-terminal conserved domains named XD ^8^. Most *Ackermannviridae* phages encode up to four different TSPs (hereafter referred to as TSP1 to TSP4). Interestingly, only two of them, TSP4 and TSP2, contain the conserved XD domains necessary for linking the four TSPs into a complex. Both TSP2 and TSP4 express an XD2 domain that interact to form the branched complex. Furthermore, TSP4 and TSP2 also express an XD3 domain interacting with TSP1 and TSP3, respectively. Finally, all four TSPs contain tandem repeat domains (TD), where the TD1 domain forms the N-terminal structural domain in TSP1 and TSP3. The role of the TD1 domains is to interact with the XD3 domains of TSP2 and TSP4. To assemble the complex, TSP4 can interact with TSP1 before or after the interaction with TSP2. In contrast, TSP3 can only interact with the XD3 domain of TSP2 following the formation of the TSP4-TSP2 complex ^8,9^. Taken together, the four TSPs of *Ackermannviridae* phages encode several conserved N-terminal domains essential to establish the TSP complex.

The host recognition of *Ackermannviridae* phages is mediated by the variable C-terminal of the four TSPs. This domain recognizes polysaccharides as receptors like lipopolysaccharide (LPS), exopolysaccharides (EPS), and capsular polysaccharides (CPS), allowing the phages to infect diverse bacteria belonging to the *Enterobacteriaceae* ^8,10–14^. We previously analyzed the TSP diversity in the *Kuttervirus*, *Agtrevirus*, *Limestonevirus*, and *Taipeivirus* genera of the *Ackermannviridae* family ^12,15^. Interestingly, a large pool of genetic diversity of receptor binding domains was revealed, suggesting that phage receptor binding proteins have been diversified to match the multitude of variations of O-antigen and K-antigens expressed by *Enterobacteriaceae* ^8,10,12,13^. While genetic variation of TSP receptor binding domains has been demonstrated, especially in the *Kuttervirus* genus, it was also observed that many of the phages express similar TSPs ^12^. For instance, 53 of the 69 *Kuttervirus* phages analyzed express a similar TSP3 that binds to *Salmonella enterica subspecies* expressing either the O4 or O9 O-antigens ^12^. In contrast, phages in the *Agtrevirus* genus express unique receptor binding proteins, meaning that only a few phages share similar TSPs ^16^. However, the high degree of conservation of the XD and TD domains may serve as sites for homologous recombination and allow for acquiring new TSPs and diversifying the branched complex of *Ackermannviridae* phages. However, this remains to be experimentally proven.

This study aims to mimic the evolution of the branched TSP complex of *Ackermannviridae* phages by investigating the acquisition of novel TSPs and their impact on the assembly and functionality of the complex. We used kuttervirus phage S117 as a model phage and showed that this phage can functionally acquire entire *tsp* genes originating from phages belonging to the *Kuttervirus* and *Agtrevirus* genera of the *Ackermannviridae* family. To expand the branched TSP complex, we designed a *tsp5* gene containing N-terminal domains interacting with the complex and the C-terminal of kuttervirus phage Det7, recognizing a novel host. Interestingly, this led to recombination between *tsp5* with *tsp4*, or deletion of *tsp3* and the truncation of *tsp4*. Our results give insight into the branched TSP network, allowing *Ackermannviridae* phages to adapt to new hosts.

## Results

### Conservation of the N-terminal TSP domains allows phage S117 to acquire entire *tsp* genes from another kuttervirus phage

The structural N-termini domains of the TSPs in *Ackermannviridae* phages are essential for assembly and, consequently, functionality of the TSP complex. At the same time, the conserved N-termini of the *tsp* genes may allow for homologous recombination between *tsp* genes, thus altering host recognition ^8,9,12^. To investigate the exchange of *tsp* genes between phages within the same genus, we used kuttervirus S117 as a model phage for acquiring novel *tsp* genes, whereas kuttervirus CBA120 was a source of such *tsp* genes. Analyzing the *tsp* gene cluster of phages S117 and CBA120 showed that *tsp1* and *tsp2* are highly similar (93.25 and 99.02% identity, respectively), whereas *tsp3* and *tsp4* only show similarity in the N-termini sequences (Figure 1A). For *tsp3*, the similarity corresponds to the TD1 and TD2 domains, whereas the anchor domain, the three XD domains, and the TD1 and TD2 of TSP4 are conserved between the two phages (Figure S1 and S2). Therefore, due to the conservation of these important N-termini domains, we hypothesized that phage S117 could acquire the entire *tsp3* or *tsp4* genes of phage CBA120 and still maintain the integrity of the TSP complex (Figure 1B). Since the C-terminal receptor binding domains of newly acquired *tsps* are different, the functionality of the TSP complex can be verified by determining the host range of the recombinant phages. For this, we used previous data showing that TSP1 and TSP3 allow S117 to infect *Salmonella enterica* subspecies Minnesota O21 and Typhimurium O4 O-antigens, respectively, whereas TSP2 binds to the O157 O-antigen of *E. coli.* The host receptor for TSP4 has yet to be identified ^12^. Similarly, TSP3 and TSP4 of phage CBA120 allow infection of *E. coli* O77 or *E. coli* O78, respectively (Plattner et al. 2019). To allow phage S117 to acquire *tsp3* or *tsp4* gene from phage CBA120, we cloned each of the *tsp3* and *tsp4* genes on a plasmid flanked with 500 bp sequences homologous to phage S117 and used a CRISPR-Cas9 system as counter selection. Homologous recombination was promoted by infecting *Salmonella* Typhimurium (LT2c ΔStyLTI), carrying the homologous template and the CRISPR-Cas9 system with phage S117. *Salmonella* Typhimurium LT2c previously deleted for the StyLTI restriction-modification system was used to maintain the two plasmids in the host ^17^. Subsequently, single plaques were picked and screened by PCR for the presence of the *tsp3* or *tsp4* genes of CBA120, respectively. Recombinant phages S117-*tsp3\** and S117-*tsp4\** were then isolated and shown to form plaques on the new hosts, *E. coli* O77 or *E. coli* O78, respectively. In addition, we confirmed that the TSP complex was still functional by showing that the recombinant phages S117-*tsp3\** and S117-*tsp4\** could still form plaques on the remaining phage S117 hosts with an efficiency of plating similar to the wildtype S117 phage (Figure 1C and D). Furthermore, sequencing the recombinant phages S117-*tsp3\** and S117-*tsp4\** confirmed the exchange of *tsp* genes. Overall, we showed that phage S117 can acquire entire *tsp3* and *tsp4* genes from a phage within the genus and still preserve the functionality of the TSP complex.

**Figure 1:**
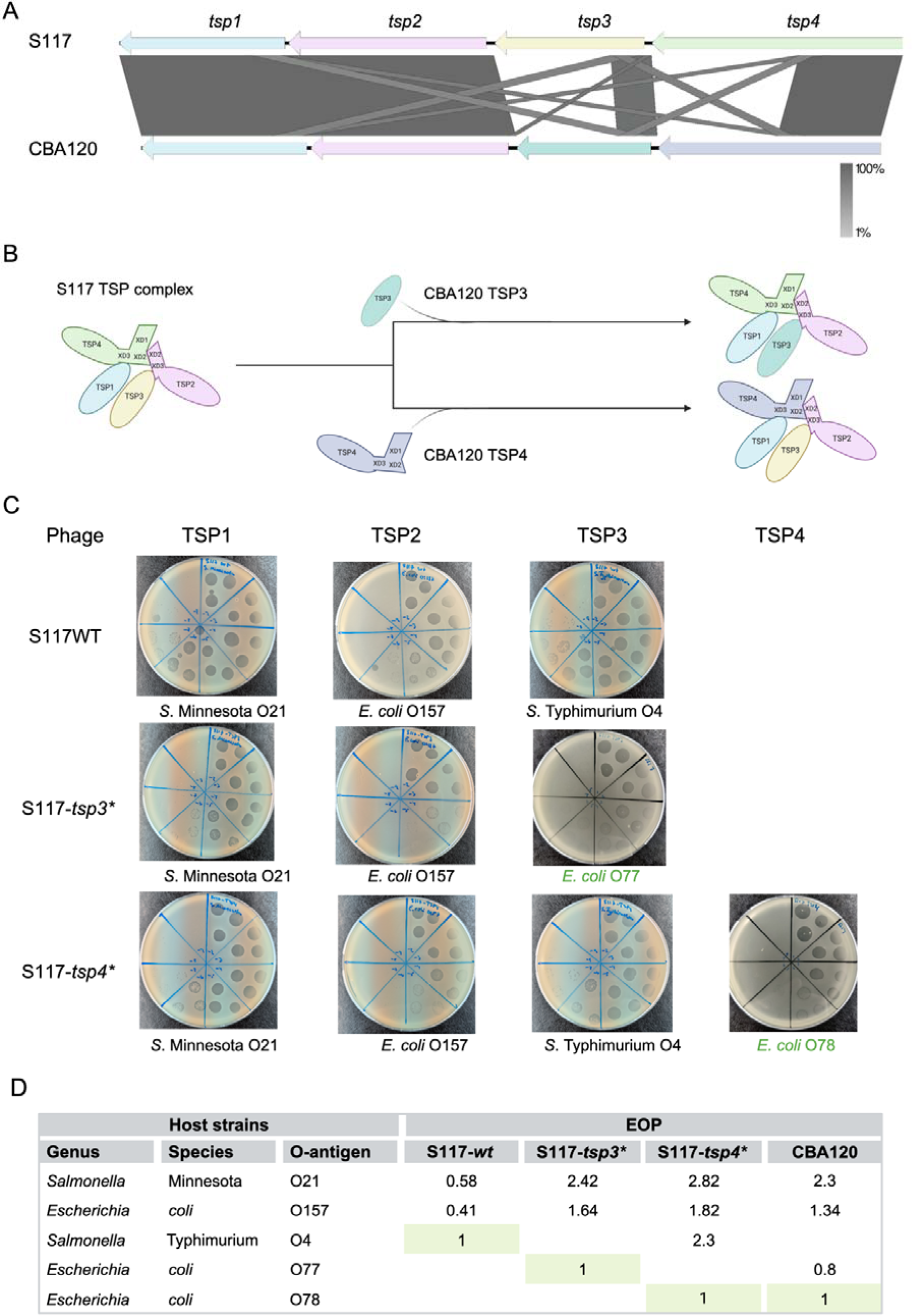
Phage S117 acquisition of *tsp3* and *tsp4* from *Kuttervirus* phage CBA120. A) Comparison of the *tsp* locus of phages CBA120 and S117 demonstrate that the phages encode similar *tsp1* and *tsp2* genes, whereas the *tsp3* and *tsp4* genes only share similarity in the N-termini. The alignment of the *tsp* gene cluster was generated by Clinker version 0.0.10 with the percentage identity between the genes indicated by the grey scale. B) Overview of the branched TSP complex of S117 before and after the acquisition of *tsp* genes from *Kuttervirus* CBA120. C) Host range analysis of S117-*wt* and recombinant phages S117-*tsp3*\* and S117-*tsp4*\* by plating 10-fold dilutions of phage stock on each host as indicated and observing plaque formation. The S117-*tsp3\** phage infects the new TSP3 host (*E. coli* O77) and the TSP1 and TSP2 hosts of S117. Phage S117-*tsp4\** infects the new TSP4 host (*E. coli* O78) and the TSP1, TSP2, and TSP3 hosts of S117. D) Efficiency of plating (EOP) of S117-*wt*, the recombinant S117-*tsp3*\* and S117-*tsp4*\* and CBA120. EOPs were calculated by dividing the PFU/mL of the tested strains by the PFU/mL of the propagation strain (indicated by light green and EOP of 1).

### Phage S117 can acquire a new *tsp2* gene from agtrevirus phage AV101

Our previous analysis of the diversity of TSPs of *Ackermannviridae* phages revealed that similar TSPs were associated with phage genera, suggesting that exchange only occurs between phages in the same genus ^12^. However, we have recently shown that the receptor binding domain of TSP4 of agtrevirus AV101 shared similarity towards kuttervirus LPST94. As a consequence, an exchange may be possible between different genera within this family ^16^. Indeed, the N-termini of *tsp* genes are highly similar, including the conserved XD domains of TSP2 and TSP4 and the TD1 domains of TSP1 and TSP3. We, therefore, speculated if *tsp* genes of agtrevirus AV101 belonging to another genus in the *Ackermannviridae* family could be exchanged by kuttervirus phage S117 and still form a functional TSP complex. Agtrevirus phage AV101 infects Extended Spectrum β-lactamase (ESBL) producing *E. coli* strains [18], and we previously showed that the four TSPs recognize *E. coli* O8, O82, O153, and O159 O-antigens, respectively ^16^. Alignment of the *tsp* gene clusters of phages AV101 and S117 show sequence similarity in the N-termini conserved domains in all four TSPs (Figure 2). For example, the XD2, XD3, and TD1 domains of TSP2 are conserved (Figure S3), suggesting that TSP2 of AV101 can be incorporated into the S117 TSP complex. To demonstrate the functionality of this recombinant TSP complex, we used the same approach described above. Still, instead of screening for the new *tsp2* gene with PCR, we directly selected for plaque formation on the new host, *E. coli* O82. The recombinant S117-*tsp2\** phage did indeed infect the new *E. coli* O82 host of AV101 TSP2 as well as the hosts of TSP1 and TSP3 of S117, *Salmonella* Minnesota O21 and *E. coli* O157, respectively (Figure 2B and C). In conclusion, these results demonstrate that a *tsp2* gene originating from *Agtrevirus* within the *Ackermannviridae* family can be exchanged across genera and still produce infectious phages.

**Figure 2:**
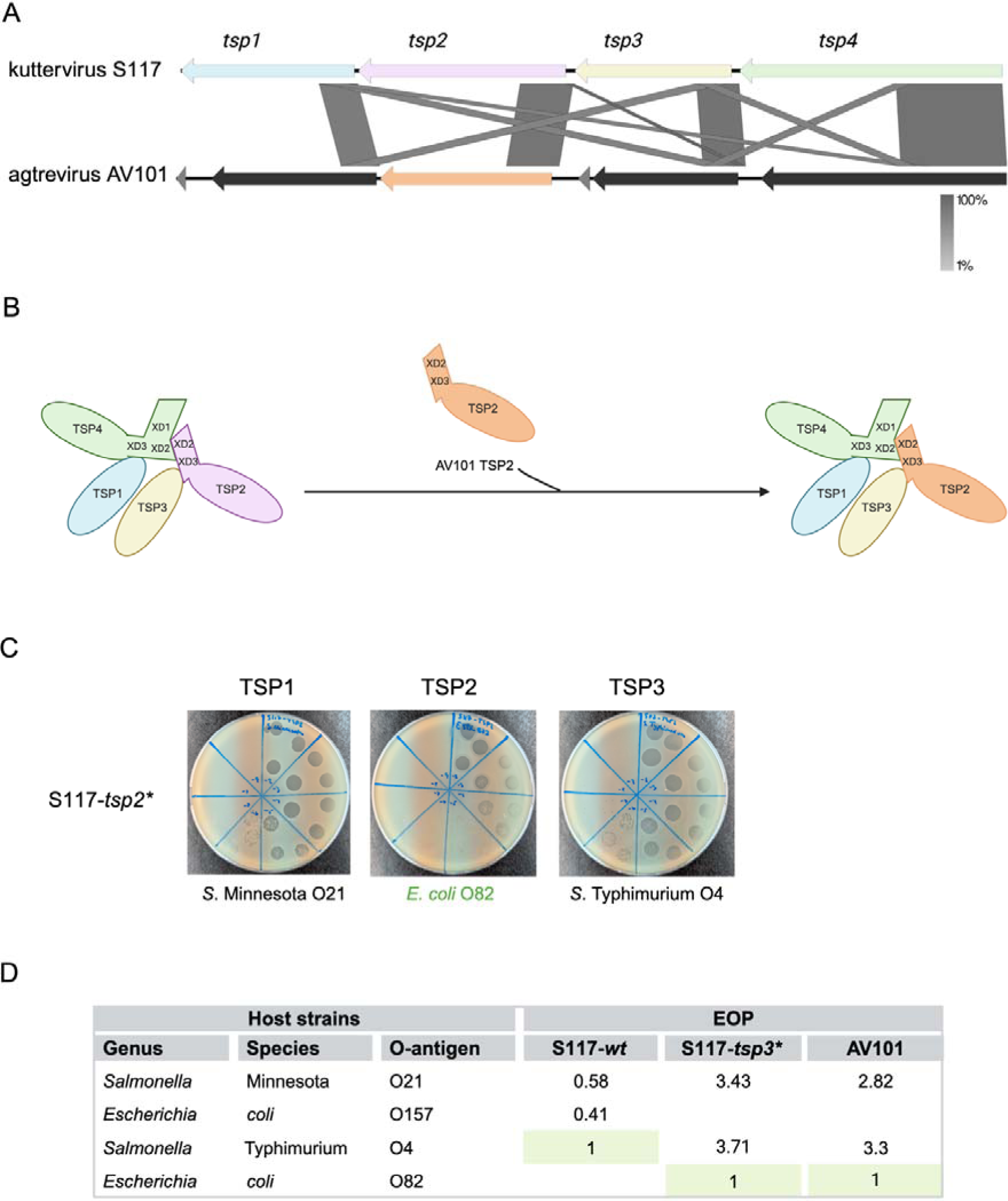
Phage S117 acquisition of *tsp2* from *Agtrevirus* phage AV101. A) Comparison of the *tsp* locus of phages AV101 and S117 showed that all genes are similar in the N-termini, and only other short regions are similar between the *tsp1*, *tsp3,* and *tsp4* genes of the two phages. The alignment of the *tsp* gene cluster was generated by Clinker version 0.0.10 with the percentage identity between the genes indicated by the grey scale. B) Overview of the branched TSP complex of S117 before and after the acquisition of *tsp2* from *Agtrevirus* phage AV101. C) Host range analysis of recombinant phage S117-*tsp2*\* by plating 10-fold dilutions of phage stock on each host as indicated and observing plaque formation. The S117-*tsp2\** phage infects the new TSP2 host (*E. coli* O82) and the TSP1 and TSP3 hosts of S117. D) Efficiency of plating (EOP) of S117-*wt*, the recombinant S117-*tsp2*\*, and AV101. EOPs were calculated by dividing the PFU/mL of the tested strains by the PFU/mL of the propagation strain (indicated by light green and EOP of 1).

### An attempt to add a *tsp5* gene led to recombination between *tsp4* and *tsp5*

The XD domains in TSP2 and TSP4 are crucial for assembling the branched TSP complex since the XD2 domains of TSP2 and TSP4 interact (Figure 3B) ^8,9^. We speculated if an additional TSP (TSP5) could be incorporated into the complex by carrying an XD2 domain, for example by addition this to an existing TSP2, as we reasoned that a TSP2 containing two XD2 domains could interact with both TSP4 and TSP2 (Figure 3B). To create such a novel *tsp5*, we added the XD2 domain of *tsp4* of S117 upstream of the entire *tsp2* gene originating from *Kuttervirus* Det7 (recognizing the O3 O-antigen from *Salmonella* Anatum) ^18^. A schematic representation of the synthetical construct is presented in Figure S4. To prevent disrupting the transcription of the remaining *tsp* genes, we aimed to insert the novel *tsp5* downstream of the *tsp* gene cluster following the successful strategy described above (Figure 3A). To isolate recombinant phages, we utilized the same methods as before and spotted the stock containing phages with potential recombinant genomes directly on the new *S.* Anatum host and isolated plaques representing S117 carrying TSP5. Indeed, a recombinant phage S117-*tsp5\** was able to infect the new *S.* Anatum host and the hosts recognized by TSP1, TSP2, and TSP3 (Figure 3C and D). As we do not know the receptor of TSP4, and hence the host of S117, we verified by PCR that all *tsp* genes, including *tsp5*, were present in the recombinant phage S117-*tsp5*\*. Our results showed that amplification of the region for the *tsp5* gene insert did not correspond to the expected size (3658 bp) but was identical to the wild type S117 (1000 bp) (Figure 3E). In addition, the *tsp4* gene of the recombinant S117-*tsp5\** genome could not amplify (Figure 3E). We hypothesized that a recombinant event between *tsp4* and *tsp5* may have created a phage able to infect the new host. To investigate this further, we sequence-verified the genome of phage S117-*tsp5*.* We found that the receptor binding domain of *tsp4* was replaced by the receptor binding domain of *tsp5*, indeed suggesting recombination between *tsp4* and *tsp5* (Figure 3F). Overall, we did not manage to introduce an additional *tsp* gene into the genome of S117 that could interact with the TSP complex. Instead, the recombination event created a phage with a *tsp4* containing the novel receptor binding domain of phage Det7, allowing the phage to infect *S*. Anatum.

**Figure 3:**
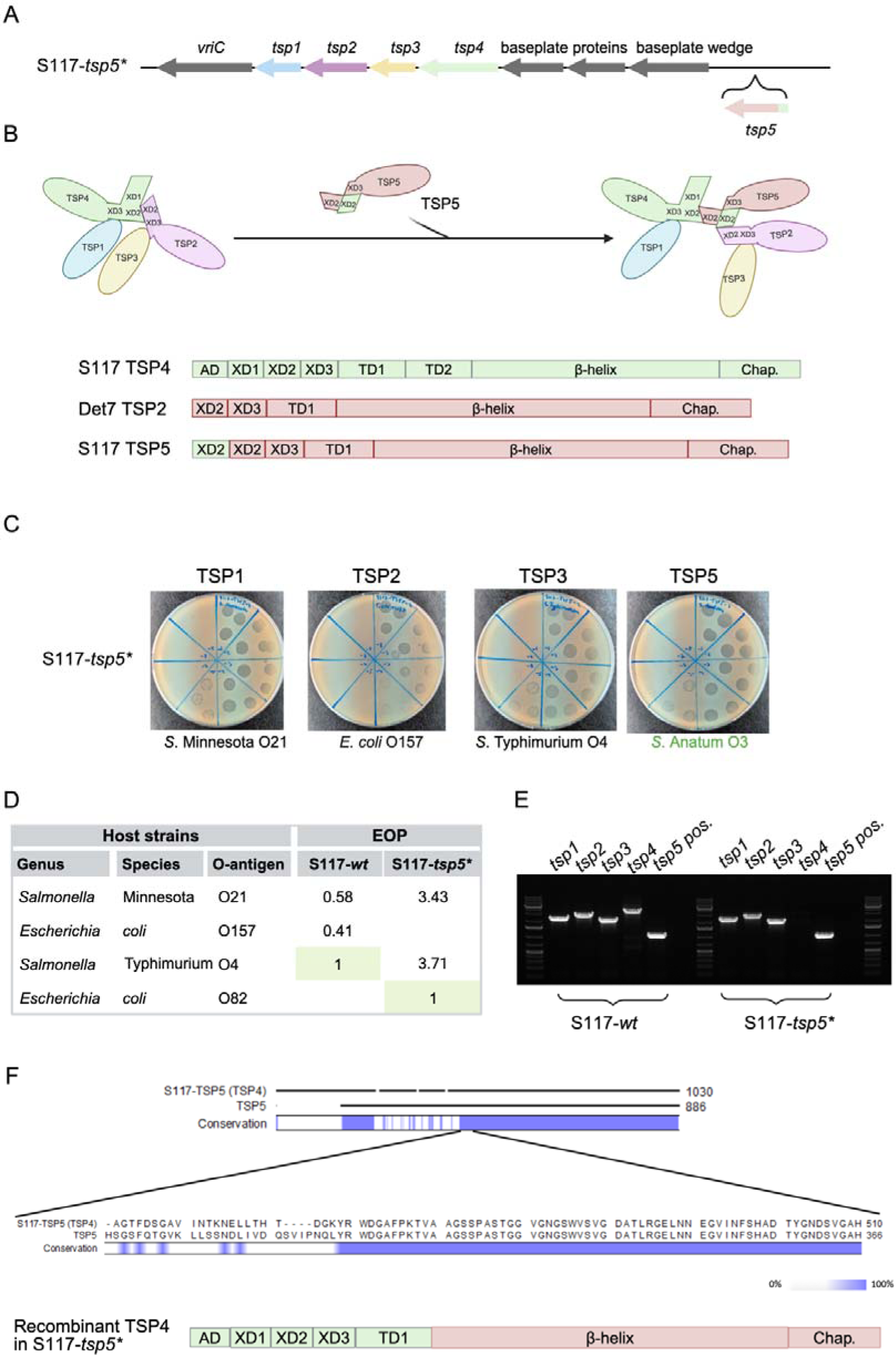
Selection for phage S117-*tsp5\** results in recombination between S117 *tsp4* and *tsp5* genes. A) Location of the additional *tsp5* gene in the genome of phage S117. B) Overview of the designing of the synthetic *tsp5* gene composed of the XD2 domain of *tsp4* of phage S117 fused to the entire *tsp2* gene of phage Det7 and model for the expanded TSP complex. C) Host range analysis of recombinant phage S117-*tsp5*\* by plating tenfold dilutions of phage stock on each host as indicated and observing plaque formation. The S117-*tsp5\** phage infects the new TSP5 host (*S*. Anatum) and the TSP1, TSP2, and TSP3 host of S117. D) Efficiency of plating (EOP) of S117-*wt* and the recombinant S117-*tsp5*\*. EOPs were calculated by dividing the PFU/mL of the tested strains by the PFU/mL of the propagation strain (indicated by light green and EOP of 1). E) PCR using primers as indicated in materials and methods to detect the four *tsp* genes and determine the location of the *tsp5* genes. F) Whole genome sequencing of recombinant S117-*tsp5*\* demonstrates a recombination event between *tsp4* and *tsp5*, resulting in a chimeric TSP4 protein containing the N-terminal of Tsp4 and the receptor binding domain of the theTSP5.

### Acquisition of a *tsp5* gene resulted in deletion of *tsp3* and truncation of *tsp4*

In a second attempt to create S117 phages carrying an additional *tsp5* gene, we used the previously isolated recombinant phage S117-*tsp4\** expressing TSP4 from CBA120 instead of wildtype S117, thus allowing for screening for all five TSP hosts (Figure 3). We indeed isolated a recombinant phage (S117-*tsp4\**-*tsp5\**) that infects *S.* Anatum (Figure 4A). Unexpectedly, phage S117-*tsp4\**-*tsp5\** could not infect hosts recognized by TSP2, TSP3, and TSP4 and could barely infect the TSP1 host (Figure 4A and B). These results suggest that the additional XD2 domain of *tsp5* may disturb the assembly of the TSP complex. Sequencing of the *tsp* cluster demonstrated that *tsp5* was correctly inserted into the genome and no reads mapped to the *tsp3* gene or the C-terminal part of the *tsp4* gene, while the recombinant phage genome could not be fully closed (Figure 4C). Yet, further analysis confirmed that the N-terminal sequences encoding the anchor domain and the XD1 and XD2 domains of the *tsp4* gene were present in our sequencing data (Figure 4D). Combining the sequencing data and host range results suggests that S117-*tsp4\**-*tsp5\** only carries the functional receptor binding domains of TSP1 and TSP5 (Figure 4E). Overall, the acquisition of a *tsp5* into the genome resulted in the deletion of *tsp3* and truncation of *tsp4* but still expressed the TSP4 domains required for assembly and functionality of the TSP complex.

**Figure 4:**
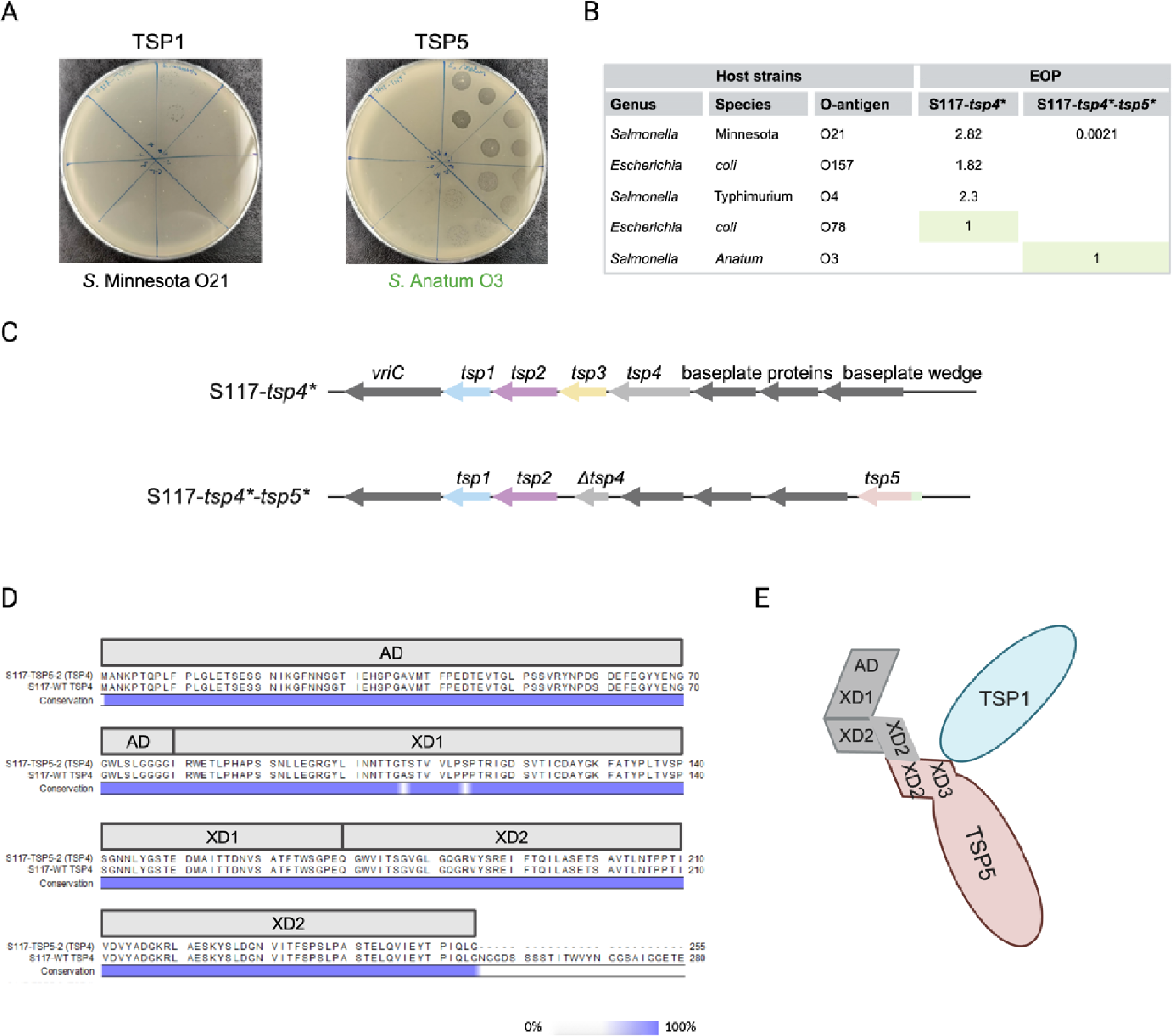
Acquisition of *tsp5* leads to deletion of *tsp3* and most of *tsp4.* A) Host range analysis of recombinant phage S117-*tsp4*\*-*tsp5*\* by plating tenfold dilutions of phage stock on each host as indicated and observing plaque formation. The new modified S117-*tsp4\**-*tsp5*\* phage infects the host of TSP5 (*S.* Anatum. B). Efficiency of plating (EOP) of S117-*wt* and the recombinant S117-*tsp4*-tsp5*\* were calculated by dividing PFU/mL of the tested strains by the PFU/mL of the propagation strain (EOP of 1). Recombinant phage S117-*tsp4\**-*tsp5*\* infects *S.* Anatum with a high EOP compared to the TSP1 host *S*. Minnesota and showed no infection of TSP2, TSP3 and TSP4 hosts. C) Whole genome sequencing of phage S117-*tsp4\**-*tsp5*\* revealed a deletion of *tsp3* and most of the *tsp4* gene. D) Sequence alignment of the N-terminal domains of TSP4 of S117-*wt* and the TSP4 of S117-*tsp4\**-*tsp5*\* showed high similarity until the XD2 domain. Only the XD domains necessary for interactions within the TSP network were conserved in the S117-*tsp4\**-*tsp5*\* phage. E) Proposed TSP complex of recombinant phage S117-*tsp4\**-*tsp5*\*.

## Discussion

Tail spike proteins (TSPs) allow phages to successfully infect their host by recognizing specific polysaccharides bacterial receptors like the LPS or CPS ^8,11,14,19^. However, bacteria express a high diversity of these polysaccharides within a species ^20–22^. For instance, *E. coli* isolates combined display an array of 185 different O-antigens of LPS ^21^. Therefore, phages expressing TSPs may be limited in their host range compared to phages that recognize a protein receptor of a more conserved nature within a bacterial species. For instance, the receptor binding protein of gelderlandvirus phage S16 recognizes the outer membrane protein OmpC as the receptor, and due to the conserved nature of the protein, the phage can infect a broad range of *Salmonella enterica* serotypes ^23^. To allow infection of a broader selection of hosts, phages using TSPs as receptor binding proteins have evolved different mechanisms to adapt to the large diversity of polysaccharide receptors. Here, we studied *Ackermannviridae* phages expressing four TSPs forming a branched complex protruding the baseplate. We aimed to mimic diverse mechanisms that allow these phages to acquire novel TSPs and investigated the impact of novel TSPs on the assembly and functionality of the complex.

Bacterial receptors are the major determinant of phage susceptibility, even more than internal phage resistance mechanisms ^1^. Hence, phages have evolved different strategies to overcome the diversity of phage receptors. One approach to adapting to a new host is to exchange receptor binding domains between phages through homologous recombination ^24–27^. Indeed, *in silico* analysis of *tsp* genes in the *Ackermannviridae* family has suggested that *tsp* genes undergo homologous recombination due to the conserved N-termini of the four genes ^12^. Our study showed that homologous recombination can exchange *tsp* genes and still produce infectious phages without disrupting the TSP complex, similar to a recent study ^28^. While we showed exchange within the family, other *in-silico* analyses of TSPs in the *Ackermannviridae* family have demonstrated that the receptor binding domains are exchanged with distant related lytic phages and prophages ^11,12,16^. For instance, the receptor binding domain of TSP4 of AV101 is similar to kayfunavirus ST31 and phapecoctavirus Ro121c4YLVW that both are distantly related phages ^16^. While domain exchange seems to be prevalent in the family, the frequency of exchange is not known. Co-evolution studies of bacteria and phages often show that phages adapt to new hosts by point mutations in the receptor binding domain and not through domain exchange ^29,30^. This may be due to the limited number of bacteria and phages in a given co-evolution experiment, thereby limiting the natural complexity. Thus, while we observe and show that exchanging whole genes or domains is a possible strategy for adaption to new hosts, the exchange frequency is unknown and remains to be determined.

While the exchange of genes of domains of receptor binding proteins is one way of adapting to a new environment, phages have also evolved to express multiple receptor binding proteins ^2–4,31^. Phages in the *Stephanstirmvirinae* subfamily, e.g. phage phi92, encode up to five receptor binding proteins including both tail spike proteins (TSPs) and tail fibers (TFs) each protruding the baseplate ^32,33^. Other phages encode proteins or domains that anchor the receptor binding proteins, like the Gp10-like domains in *Ackermannviridae* phages, to form a complex ^4,8,9^. In *Ackermannviridae* phages, it was previously suggested that the phages have evolved by gene duplication from a single TSP (TSP4) and that acquisition of the Gp10-like domains allowed multiple TSPs to interact in a complex with TSP4 ^8,12^. In most *Limestonevirus* phages, the TSP4 does not carry a receptor binding domain but only the N-terminal structural domains necessary for assembling the TSP complex, including the remaining TSPs. Furthermore, *Limestonevirus* phages do not encode a *tsp3* gene ^34–36^. Similarly, we observed that attempting to introduce a fifth TSP resulted in a recombinant phage without a TSP3 and a TSP4 with only the N-termini structural domains. Thus, TSP4 may have been an adaptor protein that established the ability to form a complex and gained a receptor binding domain through evolution. While we did not successfully add a fifth *tsp* gene in the genome of S117, two recent studies suggested that *Taipeivirus* phages like KpS110 and Menlow encode five TSPs ^4,37^. Bioinformatic analysis of these phages showed that all five genes in the *tsp* cluster encode proteins that adopt a β-helix fold common for all TSPs. Neither of the studies investigated the N-termini structural domains of the fifth TSP nor showed if the TSP is incorporated in the TSP complex. Still, overall, it may suggest that it is, in fact, possible to add a fifth TSP into the complex. In our TSP5 design, we incorporated an XD2 domain into a *tsp4* gene of Det7. The XD domains of TSP5 showed a shared sequence similarity with the XD domains present in TSP2 and TSP4 of S117. This overall sequence similarity could be the reason behind the observed different recombination events. Overall, our results also demonstrate that it is possible to engineer phages with multiple TSPs and could be used in further studies with complexes that carry Gp10-like domains.

## Method and materials

### Strains, phages, and plasmids

Bacterial strains, phages, and plasmids used in this study are found in Table 1.

**Table 1:** Bacterial strains, plasmids, and phages used in the study.

| Strains | Characteristics and use | Reference or source |
| --- | --- | --- |
| *S. Typhimurium* O4 (LT2c ΔStyLTI) | Engineering, host range | <sup>17</sup> |
| *S. Anatum* O3 | Host range | Robert Koch Institute |
| *E. coli* O157 | Host range | NCTC12900 |
| *E. coli* O77 | Host range | <sup>8</sup> |
| *E. coli* O78 | Host range | <sup>8</sup> |
| *E. coli* O82 | Host range | <sup>40</sup> |
| *E. coli* O18a | Host range | Robert Koch Institute |
| *E. coli* Stellar cells | Vector construction | Takara Bio |
| Phages |  | **Reference or source** |
| Kutternvirus CBA120 |  | <sup>8</sup> |
| Agtrevirus AV101 |  | <sup>41</sup> |
| Kutternvirus S117 wt |  | <sup>38</sup> |
| *S117-tsp2\** |  | This study |
| *S117-tsp3\** |  | This study |
| *S117-tsp4\** |  | This study |
| *S117-tsp5\** |  | This study |
| *S117-tsp4\*-tsp5\** |  | This study |
| **Plasmids** | **Resistance cassette** | **Reference or source** |
| pCas | Kanamycin | <sup>39</sup> |
| pTargetF | Spectinomycin | <sup>39</sup> |
| pTargetF-guide-TSP1 | Spectinomycin | This study |
| pTargetF-guide-TSP2 | Spectinomycin | This study |
| pTargetF-guide-TSP3 | Spectinomycin | This study |
| pTargetF-guide-TSP4 | Spectinomycin | This study |
| pTargetF-guide1-TSP5 | Spectinomycin | This study |
| pTargetF-guide2-TSP5-2 | Spectinomycin | This study |
| pTwist-Det7-TSP | Kanamycin | Twist Bioscience |

### Bioinformatic analysis

The genomes of *Kuttervirus* phages S117 (accession number MH370370) and CBA120 (NC_016570), and *Agtrevirus* phage AV101(OQ973471) were used for analysis of their *tsp* gene cluster and for designing primers for *tsp* exchange. Sequence similarities of the *tsp* gene clusters were visualized using EasyFig version 2.2.5 with default settings. Alignments of individual *tsp* genes of CBA120, AV101, and S117 were also done in CLC Workbench 22 with default settings.

### Phage propagation

Kuttervirus phages S117, CBA120, and agtrevirus phage AV101 were propagated on the host *S.* Typhimurium (LT2c), *E. coli* (NCTC12900) and *E. coli* (ESBL040) strains, respectively, as described earlier ^12^. Single colonies of the propagating strains were inoculated into LB medium (Lysogeny Broth, Merck, Darmstadt, Germany) and incubated at 37°C shaking at 170 rpm until reaching the exponential phase (approximately five hours). Previous stocks of the phages were diluted to 10^3^-10^5^ PFU/mL, and 330 µL of the phage dilutions and 330 µL of the hosts were mixed before 12mL top agar (Lbov; LB broth with 0,6% Agar bacteriological no.1, Oxoid) were added to the mixture and poured onto three LA plates (LB with 1,2% agar). After the top agar was solidified, they were incubated overnight at 37°C. The next day, 5 mL SM buffer (0.1 M NaCl, 8 mM MgSO4·7H2O, 50 mM Tris-HCl, pH 7.5) was added to each of the three plates followed by incubation of the plates overnight at 4°C and shaken at 50 rpm. The SM buffer was collected and centrifuged at 11,000 rpm at 4°C for 15 minutes and filtered twice with 0.2 µM filters. The new phage stocks were stored at 4°C before use. The titer of the stocks was determined with a plaque assay (described below).

### Phage host range analysis

To determine the host range of CBA120, AV101, and S117 phages, plaque assays were carried out ^38^. Briefly, single colonies of strains of interest were inoculated into a 5 mL medium and incubated at 37°C shaking at 180 rpm for approximately five hours to reach the exponential phase. Subsequently, 100 µL of the strain was mixed with 4 mL of top agar and poured onto LA plates. The plates were dried, and 10-fold serial dilutions of the phage stocks were spotted onto the top agar plates. After overnight incubation, the plates were screened for single plaques, and the efficiency of plating was calculated by comparing the PFU/mL of the tested strains by the PFU/mL of the propagation strain.

### Phage DNA isolation

Phages DNAs were isolated before engineering or DNA sequencing ^12^. Rnase and Dnase (Thermo Fischer Scientific) were added to the sample with a final concentration of 10 µg/mL and 20 µg/mL, respectively, prior to the nucleic acid extraction. The sample mixture was incubated for 20 min at 37°C followed by the addition of sterile EDTA (pH 8) (Thermo Fischer Scientific) at a concentration of 20 mM. To degrade the phage capsid, 50 µg/mL of Proteinase K (Thermo Fischer Scientific) was added to the sample and incubated for two hours at 56°C. The release of the DNA was verified on a 1% agarose gel. Following, the genomic DNA Clean & ConcentratorTM-10 kit (Zymo Research) was used to isolate the phage genome using the manufacturer’s instructions. The concentration of the isolated DNA was measured with Qubit (Thermo Fisher Scientific), and the quality of the DNA was verified on a 1% agarose gel.

### Phage engineering

Genetic engineering of phages was performed through CRISPR/Cas9, as previously described ^17^. Briefly, we used a recently developed two plasmids system, pEcCas (addgene #73227) and pEcgRNA (addgene #166581), where Cas9 was encoded by the pEcCas plasmid, and the Cas9 RNA guide could be cloned into the pEcgRNA plasmid as well as the recombinant template used to modify the phage genomes. Furthermore, the pEcCas plasmid also expressed the Lambda Red system under an inducible arabinose promoter to increase recombination efficiency ^39^.

### Guide RNA efficiency

Guide RNA (gRNA) targeting the four S117 TSPs were designed using the CRISPR tool on the Benchling® website. A list of the efficient guides for each tsp gene is presented in Table 2. The guides were introduced into the pEcgRNA vector through reverse PCR using the CloneAmp HiFi PCR Premix (Takara Bio) following the manufacturer’s instructions. Primers for cloning the guides into pEcgRNA are presented in the Appendix (Table S1). The PCR products were purified using Zymo PCR purification kit (Zymo Research) and directly transformed into Stellar™ chemical competent cells (Takara Bio) and plated on spectinomycin (50 µg/ml) plates. The pEcgRNA-guide plasmids from an overnight culture of Stellar cells expressing the plasmids grown in LB medium with spectinomycin (50 µg/ml) were extracted using GeneJET Plasmid Miniprep Kit (Thermo Scientific™). The pEcgRNA-guide plasmids were individually cloned into LT2c ΔStyLTI competent cells harboring the pEcCas plasmid and plated on both spectinomycin for selection of pEcgRNA (50 µg/ml) and kanamycin for selection of pEcgRNA-guide plates (50 µg/ml). The next day, the presence of both plasmids was screened by colony PCR using DreamTaq Green PCR Master Mix (Thermo Fischer™). Guide efficiency was evaluated by plaque assay on LT2c ΔStyLTI containing pEcgRNA-guide and pEcCas and on LT2c ΔStyLTI containing only pEcCas as a control (Figure S4).

**Table 2:**
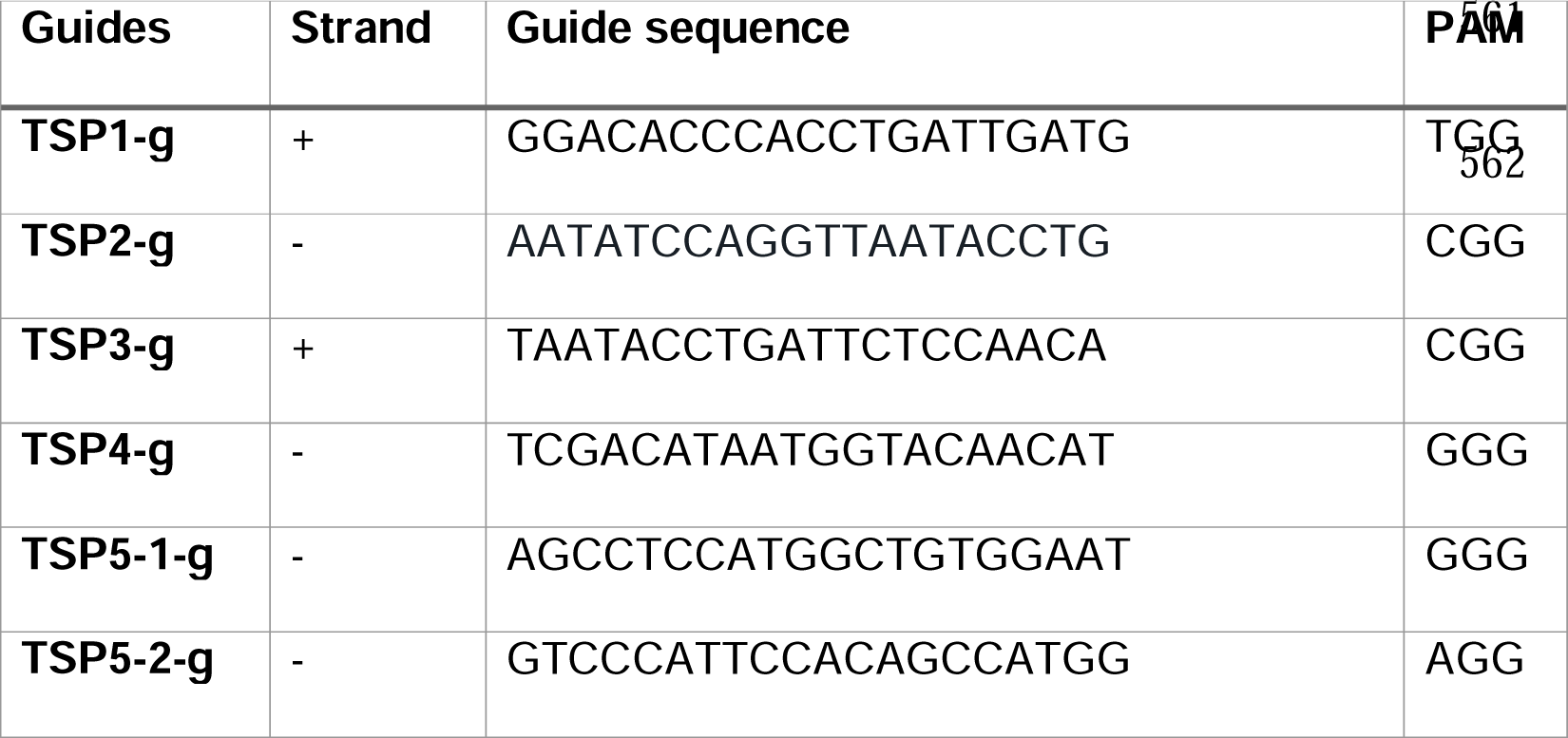
CRISPR guides used for *tsp* exchange.

### Construction of the recombinant templates

The guides with the highest efficiency were further used for *tsp* gene exchange. All primers used are presented in Table S1. To exchange the *tsp* genes in *Kuttervirus* S117, we constructed a recombinant template (RT). More specifically, the *tsp* gene and 500 base pair upstream and downstream of the S117 *tsp* gene that should be exchanged (Left and Right Homology Arms (LHA and RHA)) were amplified by PCR using CloneAmp HiFi PCR Premix (Takara Bio). LHA and RHA primers carried overhangs with homology to the *tsp* exchanged gene. After PCR purification, the RT was constructed by Splicing by Overhang Extension (SOE) PCR. Briefly, the LHA and RHA PCR products were used as primers for amplifying the *tsp* exchange gene using CloneAmp HiFi PCR Premix (Takara Bio). After five PCR cycles, primers amplifying the whole new construct were added, and the PCR was continued for 30 additional cycles. To verify the correct assembly of the three fragments 5 µL of the PCR reaction was run on a 1% agarose gel. Correct amplicons were isolated by Zymo PCR purification kit (Zymo Research). The PCR products were cloned into a PCR linearized pEcgRNA-guide plasmid to make the pEcgRNA-guide-RT plasmid using In-fusion® HD-cloning kit (Takara Bio) following the manufacturer’s instructions.

### Exchange of *tsp* genes

The LT2c expressing both pEcCas and the different pEcgRNA-guide-RT plasmids were grown to an OD600=0.2 before 0.1% arabinose was added to induce the expression of the Lambda Red system. The strains were further grown for two hours followed by a plaque assay to perform the recombination between the phage and the pEcgRNA-guide-RT plasmid. The next day, plaques were screened for the presence of the new *tsp* genes with DreamTaq Green PCR Master Mix (Thermo Fischer™). Positive plaques were then used for a new plaque assay with the new host to test if the recombinant phage could infect the host. The recombinant phages were further propagated on the new host.

### Construction of TSP5 engineered phage

To construct a phage with an additional *tsp* gene (*tsp5*) we designed a *tsp2* gene from kuttervirus phage Det7 (accession number NC_027119) with an additional XD2 domain from S117 *tsp4* gene in the start of the gene (Figure S5). Furthermore, the promoter for the *tsp2* of Det7 was added to the construct. The designed gene was ordered from Twist Bioscience. To insert the *tsp5* gene into the genome of S117 we utilized the same method as for exchanging the *tsp* genes (see Phage engineering section). Guide RNAs are presented in Table 2 and primers for cloning into pEcgRNA are presented in Table S1.

### Phage DNA sequencing

Using a Qubit 2.0 instrument and the Qubit dsDNA BR Assay Kit (Thermo Fisher Scientific), the quality of the DNA preparations of genomes of S117 and recombinant S117 was evaluated. The NEBNext Ultra DNA Library Prep kit from New England Biolabs was used to prepare the genomes for Illumina sequencing, and the HiSeq 4000 machine (Illumina) was used to sequence the samples. The CLC Genomics Workbench 21 (Qiagen, Aarhus) with default settings was used to assemble the raw reads after trimming.

## Acknowledgments

This work was supported by the Danish Council for Independent Research (9041-00159B).

## CRediT authorship contribution statement

**Anders Nørgaard Sørensen:** Conceptualization, Methodology, Validation, Formal analysis, Investigation, Writing – original draft, Writing – review & editing. **Dorottya Kalmar:** Validation, Formal analysis, Investigation. **Cedric Woudstra:** Conceptualization, Methodology, Writing – review and editing. **Jorien Poppeliers**: Formal analysis, writing - review & editing. **Rob Lavigne**: Writing - review & editing, Funding acquisition. **Martine Camilla Holst Sørensen:** Conceptualization, Writing – review & editing, Funding acquisition. **Lone Brøndsted:** Conceptualization, Project administration, Supervision, Visualization, Writing - review & editing, Funding acquisition.

## Abbreviations

CPS: Capsular polysaccharide
EPS: Exopolysaccharide
ESBL: Extended spectrum ß-lactamase
LB: Luria-Bertani
LHA: Left homology arm
LPS: Lipopolysaccharide
PFU: Plaque formation unit
RBP: Receptor binding protein
RHA: Right homology arm
RT: Recombinant template
TD: Tandem repeat
TF: Tail fiber
TSP: Tail spike protein

## References

1. Gaborieau, B., Vaysset, H., Tesson, F., Charachon, I., Dib, N., Bernier, J., Dequidt, T., Georjon, H., Clermont, O., Hersen, P., et al. (2023). Predicting phage-bacteria interactions at the strain level from genomes. bioRxiv, 2023.11.22.567924. 10.1101/2023.11.22.567924.

2. Prokhorov, N.S., Riccio, C., Zdorovenko, E.L., Shneider, M.M., Browning, C., Knirel, Y.A., Leiman, P.G., and Letarov, A. V. (2017). Function of bacteriophage G7C esterase tailspike in host cell adsorption. Mol Microbiol 105, 385–398. 10.1111/mmi.13710.

3. Ouyang, R., Costa, A.R., Cassidy, C.K., Otwinowska, A., Williams, V.C.J., Latka, A., Stansfeld, P.J., Drulis-Kawa, Z., Briers, Y., Pelt, D.M., et al. (2022). High-resolution reconstruction of a Jumbo-bacteriophage infecting capsulated bacteria using hyperbranched tail fibers. Nat Commun 13, 7241. 10.1038/s41467-022-34972-5.

4. Latka, A., Leiman, P.G., Drulis-Kawa, Z., and Briers, Y. (2019). Modeling the Architecture of Depolymerase-Containing Receptor Binding Proteins in Klebsiella Phages. Front Microbiol 10. 10.3389/fmicb.2019.02649.

5. Knecht, L.E., Veljkovic, M., and Fieseler, L. (2020). Diversity and Function of Phage Encoded Depolymerases. Front Microbiol 10, 1–16. 10.3389/fmicb.2019.02949.

6. Yap, M.L., Klose, T., Arisaka, F., Speir, J.A., Veesler, D., Fokine, A., and Rossmann, M.G. (2016). Role of bacteriophage T4 baseplate in regulating assembly and infection. Proc Natl Acad Sci U S A 113, 2654–2659. 10.1073/pnas.1601654113.

7. Taylor, N.M.I., Prokhorov, N.S., Guerrero-Ferreira, R.C., Shneider, M.M., Browning, C., Goldie, K.N., Stahlberg, H., and Leiman, P.G. (2016). Structure of the T4 baseplate and its function in triggering sheath contraction. Nature 533, 346–352. 10.1038/nature17971.

8. Plattner, M., Shneider, M.M., Arbatsky, N.P., Shashkov, A.S., Chizhov, A.O., Nazarov, S., Prokhorov, N.S., Taylor, N.M.I., Buth, S.A., Gambino, M., et al. (2019). Structure and Function of the Branched Receptor-Binding Complex of Bacteriophage CBA120. J Mol Biol 431, 3718–3739. 10.1016/j.jmb.2019.07.022.

9. Chao, K.L., Shang, X., Greenfield, J., Linden, S.B., Alreja, A.B., Nelson, D.C., and Herzberg, O. (2022). Structure of Escherichia coli O157:H7 bacteriophage CBA120 tailspike protein 4 baseplate anchor and tailspike assembly domains (TSP4-N). Sci Rep 12. 10.1038/s41598-022-06073-2.

10. Kabanova, A.P., Shneider, M.M., Korzhenkov, A.A., Bugaeva, E.N., Miroshnikov, K.K., Zdorovenko, E.L., Kulikov, E.E., Toschakov, S. V., Ignatov, A.N., Knirel, Y.A., et al. (2019). Host specificity of the dickeya bacteriophage PP35 is directed by a tail spike interaction with bacterial o-antigen, enabling the infection of alternative non-pathogenic bacterial host. Front Microbiol 10, 1–11. 10.3389/fmicb.2018.03288.

11. Walter, M., Fiedler, C., Grassl, R., Biebl, M., Rachel, R., Hermo-Parrado, X.L., Llamas-Saiz, A.L., Seckler, R., Miller, S., and van Raaij, M.J. (2008). Structure of the Receptor-Binding Protein of Bacteriophage Det7: a Podoviral Tail Spike in a Myovirus. J Virol 82, 2265–2273. 10.1128/jvi.01641-07.

12. Sørensen, A.N., Woudstra, C., Sørensen, M.C.H., and Brøndsted, L. (2021). Subtypes of tail spike proteins predicts the host range of Ackermannviridae phages. Comput Struct Biotechnol J 19. 10.1016/j.csbj.2021.08.030.

13. Hsu, C.R., Lin, T.L., Pan, Y.J., Hsieh, P.F., and Wang, J.T. (2013). Isolation of a Bacteriophage Specific for a New Capsular Type of Klebsiella pneumoniae and Characterization of Its Polysaccharide Depolymerase. PLoS One 8. 10.1371/journal.pone.0070092.

14. Oliveira, H., Pinto, G., Mendes, B., Dias, O., Hendrix, H., Akturk, E., Noben, J.-P., Gawor, J., Łobocka, M., Lavigne, R., et al. (2020). Tailspike with EPS-depolymerase activity, encoded by a phage belonging to a new genus, makes Providencia stuartii susceptible to serum-mediated killing. Appl Environ Microbiol. 10.1128/aem.00073-20.

15. Sørensen, A.N., Kalmar, D., Lutz, V.T., Klein-Sousa, V., Taylor, N.M.I., Sørensen, M.C.H., and Brøndsted, L. (2023). Agtrevirus phage AV101 recognizes four different O-antigens infecting diverse E. coli. microLife. 10.1093/femsml/uqad047.

16. Sørensen, A.N., Kalmar, D., Lutz, V.T., Klein-Sousa, V., Taylor, N.M.I., Sørensen, M.C.H., and Brøndsted, L. (2023). Agtrevirus phage AV101 infect diverse extended spectrum β-lactamase E. coli by recognizing four different O-antigens. bioRxiv, 2023.09.19.558411. 10.1101/2023.09.19.558411.

17. Woudstra, C., Sørensen, A.N., and Brøndsted, L. (2023). Engineering of Salmonella Phages into Novel Antimicrobial Tailocins. Cells 12, 2637. 10.3390/cells12222637.

18. Broeker, N.K., Roske, Y., Valleriani, A., Stephan, M.S., Andres, D., Koetz, J., Heinemann, U., and Barbirz, S. (2019). Time-resolved DNA release from an O-antigen–specific Salmonella bacteriophage with a contractile tail. Journal of Biological Chemistry 294, 11751–11761. 10.1074/jbc.RA119.008133.

19. Domingo-Calap, P., Beamud, B., Mora-Quilis, L., González-Candelas, F., and Sanjuán, R. (2020). Isolation and characterization of two klebsiella pneumoniae phages encoding divergent depolymerases. Int J Mol Sci 21. 10.3390/ijms21093160.

20. Reeves, P.R., Cunneen, M.M., Liu, B., and Wang, L. (2013). Genetics and Evolution of the Salmonella Galactose-Initiated Set of O Antigens. PLoS One 8, 1–22. 10.1371/journal.pone.0069306.

21. Liu, B., Furevi, A., Perepelov, A. V, Guo, X., Cao, H., Wang, Q., Reeves, P.R., Knirel, Y.A., Wang, L., and Widmalm, G. (2020). Structure and genetics of *Escherichia coli* O antigens. FEMS Microbiol Rev 44, 655–683. 10.1093/femsre/fuz028.

22. Follador, R., Heinz, E., Wyres, K.L., Ellington, M.J., Kowarik, M., Holt, K.E., and Thomson, N.R. (2016). The diversity of Klebsiella pneumoniae surface polysaccharides. Microb Genom 2, e000073. 10.1099/mgen.0.000073.

23. Marti, R., Zurfluh, K., Hagens, S., Pianezzi, J., Klumpp, J., and Loessner, M.J. (2013). Long tail fibres of the novel broad-host-range T-even bacteriophage S16 specifically recognize Salmonella OmpC. Mol Microbiol 87, 818–834. 10.1111/mmi.12134.

24. Haggard-Ljungquist, E., Halling, C., and Calendar, R. (1992). DNA Sequences of the Tail Fiber Genes of Bacteriophage P2: Evidence for Horizontal Transfer of Tail Fiber Genes among Unrelated Bacteriophages We have determined the DNA sequence of the bacteriophage P2 tail genes G and H, which code for.

25. Te, F., Tart, Â., Desplats, C., and Krisch, H.M. (1998). Genome Plasticity in the Distal Tail Fiber Locus of the T-even Bacteriophage: Recombination between Conserved Motifs Swaps Adhesin Specificity.

26. Barbirz, S., Müller, J.J., Uetrecht, C., Clark, A.J., Heinemann, U., and Seckler, R. (2008). Crystal structure of Escherichia coli phage HK620 tailspike: Podoviral tailspike endoglycosidase modules are evolutionarily related. Mol Microbiol 69, 303–316. 10.1111/j.1365-2958.2008.06311.x.

27. Broeker, N.K., and Barbirz, S. (2017). Not a barrier but a key: How bacteriophages exploit host’s O-antigen as an essential receptor to initiate infection. Mol Microbiol 105, 353–357. 10.1111/mmi.13729.

28. Gil, J., Paulson, J., Brown, M., Zahn, H., Nguyen, M.M., Eisenberg, M., and Erickson, S. (2023). Tailoring the Host Range of Ackermannviridae Bacteriophages through Chimeric Tailspike Proteins. Viruses 15, 286. 10.3390/v15020286.

29. Subramanian, S., Dover, J.A., Parent, K.N., and Doore, S.M. (2022). Host Range Expansion of Shigella Phage Sf6 Evolves through Point Mutations in the Tailspike. J Virol 96. 10.1128/jvi.00929-22.

30. Boon, M., Holtappels, D., Lood, C., van Noort, V., and Lavigne, R. (2020). Host Range Expansion of Pseudomonas Virus LUZ7 Is Driven by a Conserved Tail Fiber Mutation. Phage 1, 87–90. 10.1089/phage.2020.0006.

31. Gebhart, D., Williams, S.R., and Scholl, D. (2017). Bacteriophage SP6 encodes a second tailspike protein that recognizes Salmonella enterica serogroups C2 and C3. Virology 507, 263–266. 10.1016/j.virol.2017.02.025.

32. Schwarzer, D., Buettner, F.F.R., Browning, C., Nazarov, S., Rabsch, W., Bethe, A., Oberbeck, A., Bowman, V.D., Stummeyer, K., Muhlenhoff, M., et al. (2012). A Multivalent Adsorption Apparatus Explains the Broad Host Range of Phage phi92: a Comprehensive Genomic and Structural Analysis. J Virol 86, 10384–10398. 10.1128/jvi.00801-12.

33. Schwarzer, D., Browning, C., Stummeyer, K., Oberbeck, A., Mühlenhoff, M., Gerardy-Schahn, R., and Leiman, P.G. (2015). Structure and biochemical characterization of bacteriophage phi92 endosialidase. Virology 477, 133–143. 10.1016/j.virol.2014.11.002.

34. Adriaenssens, E.M., van Vaerenbergh, J., Vandenheuvel, D., Dunon, V., Ceyssens, P.J., de Proft, M., Kropinski, A.M., Noben, J.P., Maes, M., and Lavigne, R. (2012). T4-related bacteriophage LIMEstone isolates for the control of soft rot on potato caused by “Dickeya solani.” PLoS One 7. 10.1371/journal.pone.0033227.

35. Day, A., Ahn, J., and Salmond, G.P.C. (2018). Jumbo bacteriophages are represented within an increasing diversity of environmental viruses infecting the emerging phytopathogen, Dickeya solani. Front Microbiol 9, 1–15. 10.3389/fmicb.2018.02169.

36. Day, A., Ahn, J., Fang, X., and Salmond, G.P.C. (2017). Environmental bacteriophages of the emerging enterobacterial phytopathogen, dickeya solani, show genomic conservation and capacity for horizontal gene transfer between their bacterial hosts. Front Microbiol 8, 1–9. 10.3389/fmicb.2017.01654.

37. Volozhantsev, N. V., Verevkin, V. V., Krasilnikova, V.M., Kislichkina, A.A., and Popova, A. V. (2023). Complete Genome Sequence of Klebsiella pneumoniae Bacteriophage KpS110, Encoding Five Tail-Associated Proteins with Putative Polysaccharide Depolymerase Domains. Microbiol Resour Announc 12. 10.1128/mra.00153-23.

38. Gencay, Y.E., Gambino, M., Prüssing, T.F., and Brøndsted, L. (2019). The genera of bacteriophages and their receptors are the major determinants of host range. Environ Microbiol 21, 2095–2111. 10.1111/1462-2920.14597.

39. Li, Q., Sun, B., Chen, J., Zhang, Y., Jiang, Y., and Yang, S. (2021). A modified pCas/pTargetF system for CRISPR-Cas9-assisted genome editing in Escherichia coli. Acta Biochim Biophys Sin (Shanghai) 53, 620–627. 10.1093/abbs/gmab036.

40. Vitt, A.R., Sørensen, M.C.H., Bortolaia, V., and Brøndsted, L. (2023). A Representative Collection of Commensal Extended-Spectrum-and AmpC-β-Lactamase-Producing Escherichia coli of Animal Origin for Phage Sensitivity Studies. PHAGE: Therapy, Applications, and Research 4, 35–45. 10.1089/phage.2023.0002.

41. Vitt, A.R., Sørensen, A.N., Bojer, M.S., Bortolaia, V., Sørensen, M.C.H., and Brøndsted, L. (2023). A collection of diverse bacteriophages for biocontrol of ESBL-and AmpC-β-lactamase-producing E. coli. bioRxiv, 2023.09.14.557699. 10.1101/2023.09.14.557699.

