## Supplementary information for "The branched receptor binding complex of *Ackermannviridae* phages promotes adaptative host recognition"

### Supplementary data

**Table S1: List of primers for *tsp* exchange:**

| Primers | Sequences (5’-3’) |
| --- | --- |
| TSP2: |  |
| S117_TSP2_LHA_F | CTATTACCCTGTTATCGCGCCAAGACCAATACCAC |
| S117_TSP2_LHA_R | AAGATTAGTAATTAAAAATATACCCCTCTTCGGAGGGG |
| AV101_TSP2_F | AAGAGGGGTATATTTTTAATTACTAATCTTACCACGTATGGATAGCTCA |
| AV101_TSP2_R | ATGGGGTATTTTCAAATGACCAGAAACGTCGAGAGC |
| S117_TSP2_RHA_F | GACGTTTCTGGTCATTTGAAAATACCCCATAAAGGATTTGCCA |
| S117_TSP2_RHA_R | CATGAACTCGAGTAGGTTACTATGTACCTATATAAGCGATATTCAGGGAG |
| TSP3: |  |
| S117_TSP3_LHA_F | TATTACCCTGTTATCCCTACTCGACGGGTGTTTTGTACTCGCCACC |
| S117_TSP3_LHA_R | TGGAGCTGATGTTTGATTTATCTTGAGCAAAAACCCCGCT |
| CBA120_TSP3_F | TTTTGCTCAAGATAAATCAAACATCAGCTCCAGCCG |
| CBA120_TSP3_R | ATCCTTTATGGGGTATTATCATGATTTCTCAATTCAATCAACCACGCG |
| S117_TSP3_RHA_F | TGATTGAATTGAGAAATCATGATAATACCCCATAAAGGATGACCAATATGGG |
| S117_TSP3_RHA_R | TTATGGAGCTGCACATGAACGGTGTTGTGCAATTCACTGCTGATATTAAAAC |
| TSP4: |  |
| S117_TSP4_LHA_F | CTATTACCCTGTTATCCAACTTTAGGTAATGCACCATCCCATC |
| S117_TSP4_LHA_R | TTACGGATTGATATGATAAAAACCCCGCTTCGGCG |
| CBA120_TSP4_F | CGAAGCGGGGTTTTTATCATATCAATCCGTAAATGACATTGTGTATTGC |
| CBA120_TSP4_R | TTTAGGGGTATTACAAATGGCCAACAAACCAACACAGC |
| S117_TSP4_RHA_F | TTGGTTTGTTGGCCATTTGTAATACCCCTAAATGTATTCATGTCATCTAGG |
| S117_TSP4_RHA_R | CATGAACTCGAGTAGGTAAAGTCGAGGTCAGCACATATAATACGAA |
| TSP5: |  |
| S117_TSP5_LHA_F | TATTACCCTGTTATCTCCATGATGATCTCATTGGGGGC |
| S117_TSP5_LHA_R | GTTGATTTTAACTAATGGCTGTGGAATGGGACTGC |
| Det7_TSP5_F | CCCATTCCACAGCCATTAGTTAAAATCAACAAAACCTTTTAATATAAACGATGAT |
| Det7_TSP5_R | TGTGAGTGTTAACGATAAGTAACTTATGCCCCGCTTTGG |
| S117_TSP5_RHA_F | GGCATAAGTTACTTATCGTTAACACTCACAACTCGAAGG |
| S117_TSP5_RHA_R | ATGAACTCGAGTAGGCAATTTGATTCCAACGCGGGTG |

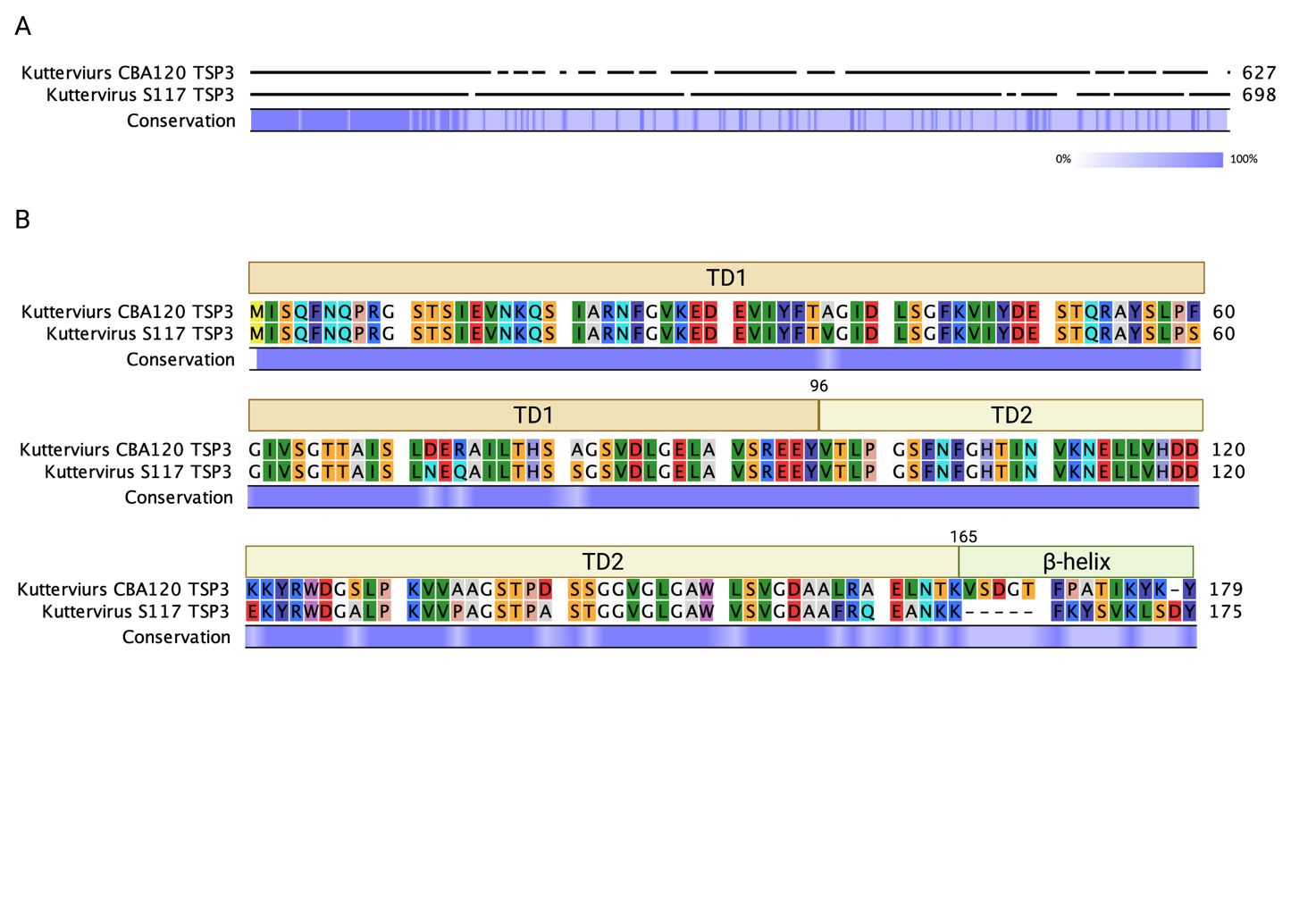
**Figure S1: *tsp3* sequence similarities between kuttervirus phages S117 and CBA120.** A) Alignment of the two genes showed that only the N-termini sequences are similar. B) Zoom in on the N-termini alignment. The structural domains are shown to indicate what the sequence encode for in the TSP.

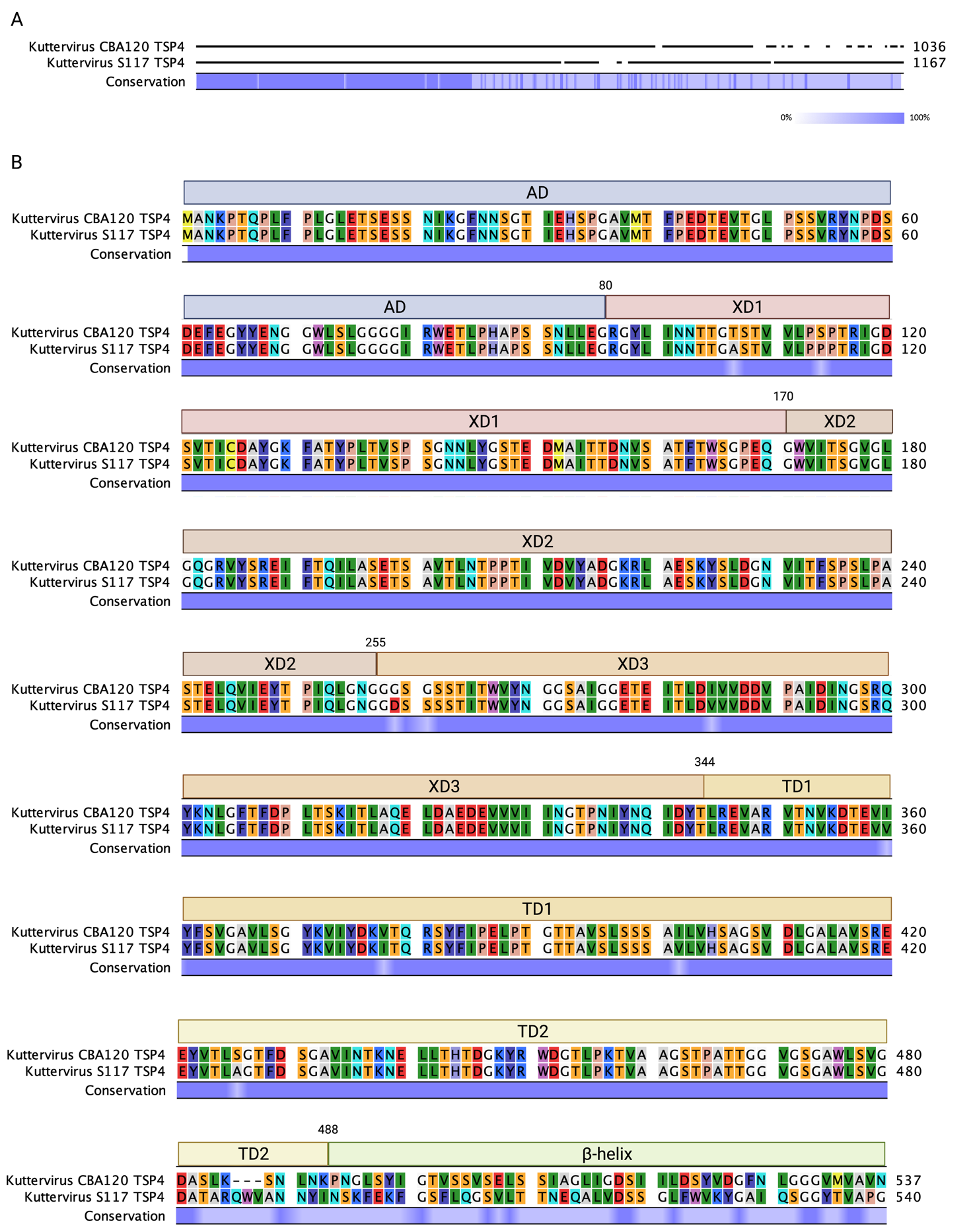

**Figure S2: Sequence alignment of the *tsp4* genes of kuttervirus phages CBA120 and S117**. A) Similarity over the entire length of the two genes. B) The sequences encoding N-termini structural domains are conserved between the two genes.

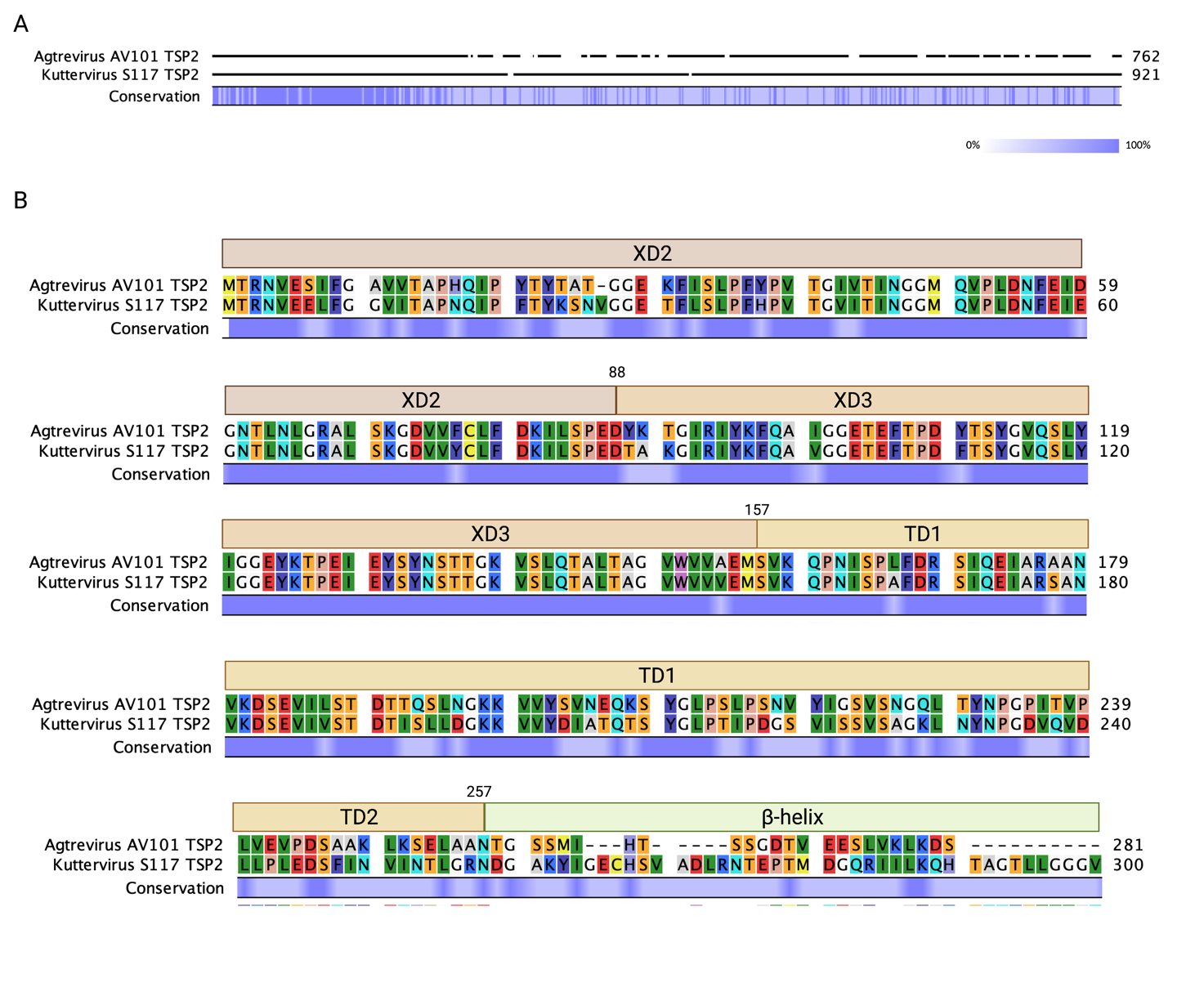

**Figure S3: The tsp2 genes from agtrevirus phage AV101 and kuttervirus phage S117 are similar in the N-termini.** A) Alignment of the tsp2 genes. B) Zoom in of the N-termini showed that the N-termini of the two genes are highly similar.

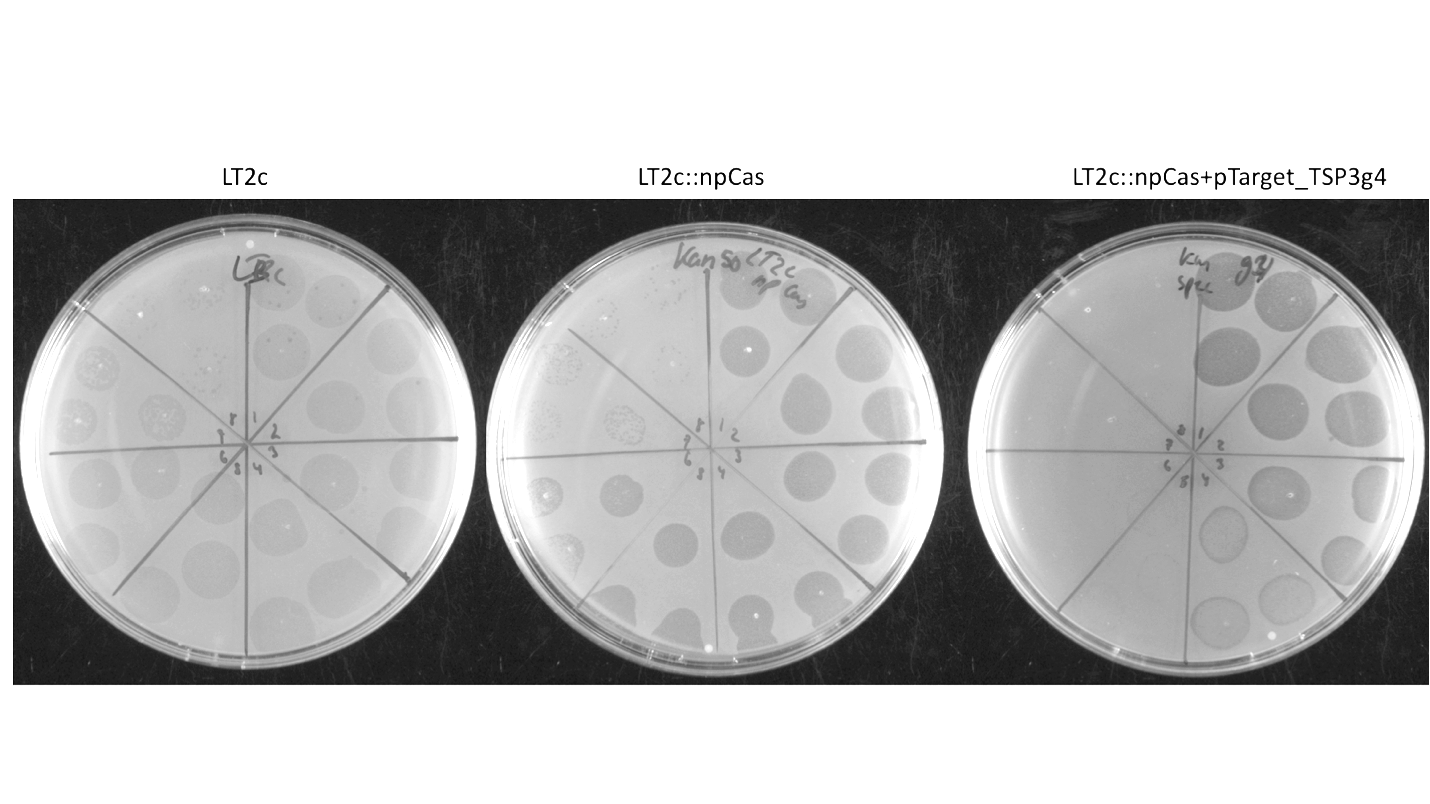

**Figure S4: Evaluation of the guide efficiency**. The efficiency of the guides was checked by spotting S117 on LT2c with pEcCas and pEcgRNA-guides. LT2c and LT2c with pEcCas were used as controls. Reduced efficiency of plating demonstrates that the guide is effective.

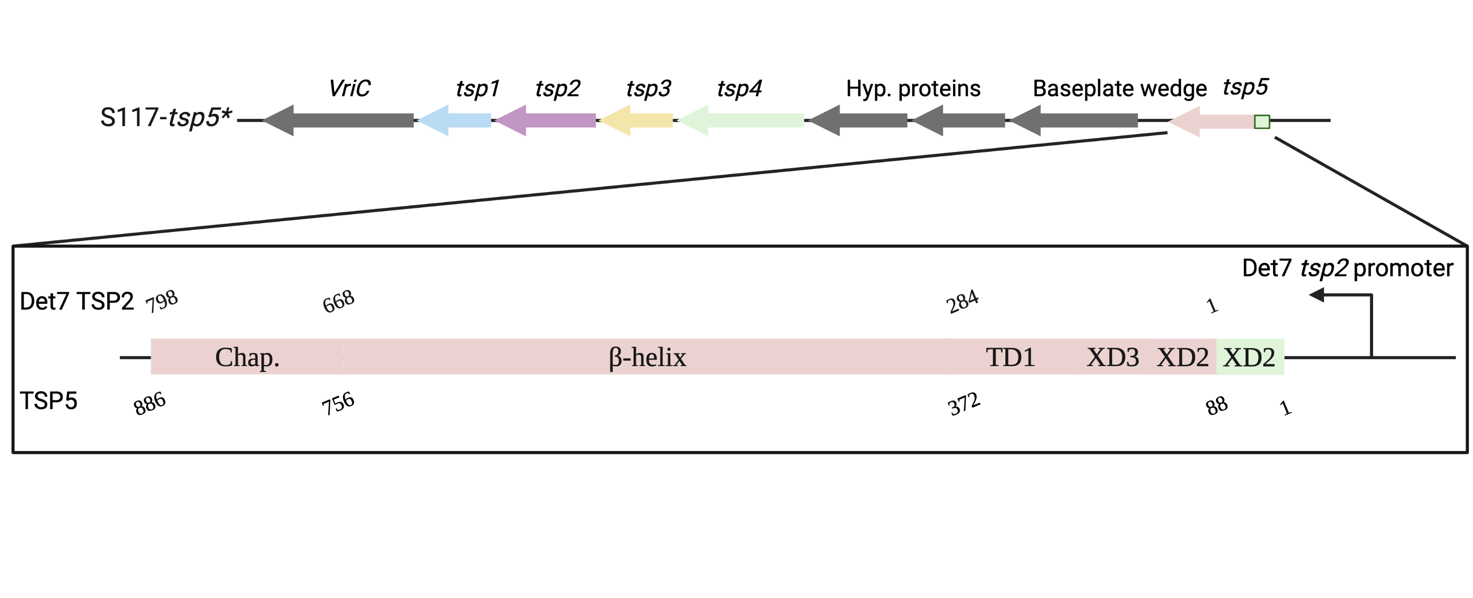

**Figure S5: Construction of the tsp5.** Tsp5 was inserted downstream of the tsp gene cluster. The TSP5 was made up of TSP2 from kuttervirus Det7 and an XD2 domain from kuttervirus S117 was added in front of the TSP2. The promoter for tsp2 from kuttervirus Det7 was also added to the construct.
